# Stable Network Homeostasis during Multi-Level Postnatal Maturation of the Mouse Tuberoinfundibular Dopamine–Prolactin Axis

**DOI:** 10.64898/2026.07.16.737908

**Authors:** Andrea Locarno, Andriana Mantzafou, Jimena Ferraris, Christian Broberger

**Affiliations:** Department of Biochemistry and Biophysics, Stockholm University, SE-106 91 Stockholm, Sweden

## Abstract

The hypothalamus orchestrates endocrine function via specialized neuronal populations that interface with the pituitary gland. While postnatal circuit refinement is a hallmark of most neural systems, its contribution to hypothalamic neuroendocrine networks remains elusive. Among these populations, tuberoinfundibular dopamine (TIDA) neurons of the arcuate nucleus are the primary source of tonic inhibition of prolactin (Prl), a hormone essential for reproduction, parental physiology, and behavior. While Prl levels surge during early mouse postnatal life, it remains unclear whether the TIDA system is fully developed at birth or undergoes functional maturation.

Here, we combined immunofluorescence, slice electrophysiology, and Ca^2+^ imaging to determine the development of TIDA neurons in mice during the first three postnatal weeks. Expression of dopaminergic markers was sparse at birth but rose substantially after the first week, followed by the onset of median eminence innervation by TIDA axons. Moreover, TIDA firing rate and oscillation frequency progressively increased with age, with action potential properties and rhythmicity maturing in tandem. Strikingly, local network parameters remained stable despite ongoing changes in single-cell properties, as did excitation/inhibition (E/I) balance. These neuronal adaptations paralleled a significant rise in circulating Prl levels.

Together, our results delineate a multi-level developmental program within the TIDA system, encompassing molecular, electrophysiological, and endocrine changes. This maturation likely underlies the emergence of functional hypothalamic control over Prl secretion in early life and points to a coördinated controller-effector co-development. Our findings highlight a critical window during which TIDA neuron plasticity may influence long-term neuroendocrine function and reproductive behavior.

**Significance Statement:** Understanding how neuroendocrine circuits mature is critical for elucidating the developmental origins of hormonal homeostatic control. We reveal that tuberoinfundibular dopamine (TIDA) neurons, which control reproduction by inhibiting the release of the pituitary hormone, prolactin (Prl), undergo a tightly orchestrated postnatal maturation program encompassing population expansion, electrophysiological refinement, and synaptic scaling. Surprisingly, although individual cellular features undergo substantial shifts, network behavior remains remarkably consistent, in parallel with a gradual rise in Prl. This multi-level developmental sequence likely enables the gradual assembly of a functional TIDA-Prl axis. By defining the timeline of TIDA circuit maturation, our work provides a framework for exploring how early-life perturbations may disrupt neuroendocrine development and lead to lifelong alterations in reproductive and parental behaviors.

## Introduction

Hormones originating from the pituitary gland regulate fundamental survival functions such as reproduction, growth, stress, metabolism, and fluid balance, and are maintained within precisely controlled intervals, which can be adapted to changing circumstances of life. This exact regulation is accomplished through master control from neuroendocrine neurons of the hypothalamus (1), whose activity relies critically on feedback from the target hormone. While the mechanisms of such feedback have been the subject of intense scrutiny, one issue that has largely escaped experimental attention is to what extent the properties of the hypothalamic master control are fixed from birth or undergo postnatal maturation as adult hormone levels are established.

Many mammalian species are born in an underdeveloped state, with immature sensory and motor systems, and are dependent on parental care for their survival. In such altricial species, early postnatal life represents a continuation of central nervous system development, during which neural circuits undergo structural remodeling and functional refinement (*e.g*., 2, 3; see also 4). This period is critical for the maturation of sensorimotor integration, behavioral responses, and homeostatic regulation (5–7).

This developmental window is increasingly recognized in neuroendocrine systems, where hypothalamic-pituitary axes remodel in ways that shape lifelong physiology and behavior (see 8). For instance, earlylife modulation of the hypothalamic-pituitary-adrenal axis produces lasting effects on stress reactivity (9), while the maturation of gonadotropin-releasing hormone and kisspeptin-producing neurons plays a key role in puberty onset (10–12; see 13).

As with stress and reproductive axes, the prolactin (Prl) system also undergoes marked developmental remodeling. In adulthood, Prl’s role extends well beyond lactation, encompassing reproduction, female and male parental behaviors, and immune regulation (*e.g*., 14–18; see also 19). Establishing appropriate Prl control during development is therefore essential for programming adult homeostasis and behavior.

Importantly, the Prl axis undergoes major developmental changes after birth. The Prl-secreting cells— lactotrophs—in the anterior pituitary expand dramatically in the early postnatal period (20–22), paralleled by a marked increase in circulating Prl levels (23–25). These findings suggest the intriguing possibility that the regulatory architecture of the Prl axis is assembled progressively, rather than being fully established at birth.

The principal regulator of Prl secretion is a population of dopaminergic cells in the dorsomedial arcuate nucleus (dmArc) of the hypothalamus, known as tuberoinfundibular dopamine (TIDA) neurons. These neurons release dopamine into the hypophyseal portal system, providing tonic inhibition of lactotrophs (see 26, 27), constraining the secretion of Prl to lactation and some other distinct physiological states. Prl in turn reciprocally activates TIDA neurons via Prl receptor signaling, constituting a negative feedback loop (28, 29). In adult rodents, TIDA neurons can exhibit rhythmic oscillatory firing patterns functionally linked to Prl regulation. In male rats, this activity is a near-universal feature (30), whereas in mice, only a subset of neurons oscillates, a species difference that has been traced to the existence and absence, respectively, of electrical synapses connecting these cells (31). Optogenetic manipulations of these oscillations alter circulating Prl levels, establishing a causal link between TIDA rhythmicity and hormone secretion (18). Oscillatory dynamics are a hallmark of neuroendocrine function (see 32), and in TIDA neurons, they fine-tune dopamine release at the neurohemal interface in the median eminence (ME; 33, 34). However, how these oscillatory dynamics emerge and mature postnatally remains unknown.

The maturation of TIDA neurons likely involves anatomical and transcriptional changes as well as adaptations in excitability, synaptic integration, and network connectivity. Other hypothalamic circuits provide precedents, underscoring that functional specialization often emerges only after birth. Projections from feeding-regulatory arcuate nucleus neurons are not fully established until the second postnatal week, highlighting a delayed maturation of output connectivity (35, 36). This gradual postnatal axonal arborization parallels a transition from suckling to adult feeding behavior (37–39 These findings suggest that hypothalamic circuits are not functionally mature at birth, but instead gradually acquire their adult-like properties.

Here, we combined anatomical, electrophysiological, and imaging approaches to investigate the developmental trajectory of TIDA neurons during the early postnatal period. We identify a substantial expansion of the dopaminergic dmArc population accompanied by coördinated functional maturation that provides a model for the emergence of TIDA neurons as key regulators of circulating Prl.

## Results

### The dmArc dopaminergic population expands postnatally

Circulating Prl increases markedly during the early postnatal period in rodents (23– In agreement with these reports, we observed a significant age-dependent increase in the levels of circulating Prl across the first three postnatal weeks (**Fig. 1A**; \**p* < 0.05).

**Figure 1.**
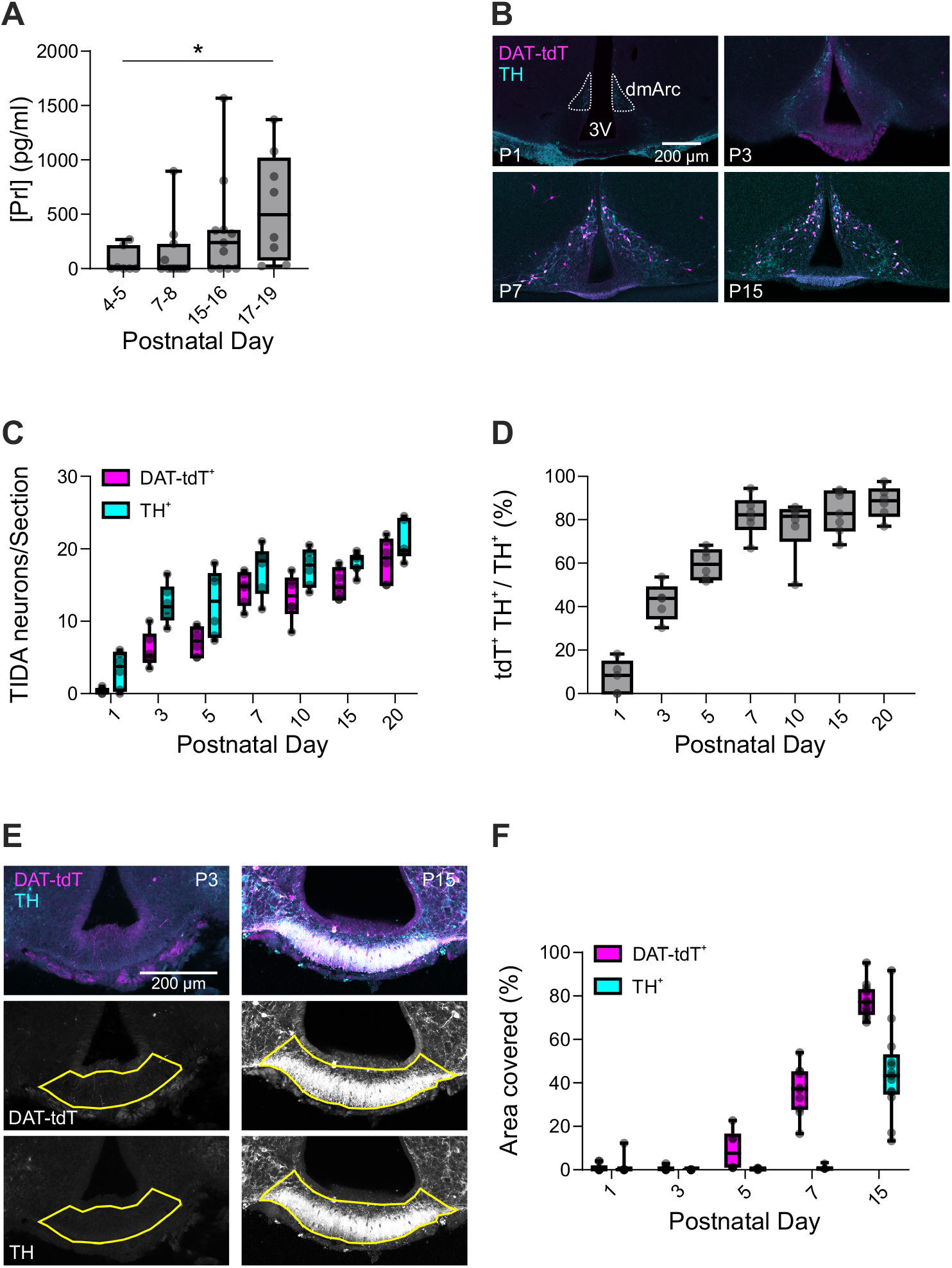
Postnatal increase in circulating Prl and expansion of the dmArc dopaminergic population. **(A)** Serum Prl concentrations across post-natal development (n = 7– 11 mice; Kruskal–Wallis test: *H*(3) = 9.23, ^*^*p* = 0.03). **(B)** Representative confocal images of the mediobasal hypothalamus at P1, P3, P7, and P15 showing DAT-tdT^+^ (magenta) and TH^+^ (cyan) neurons. Scale bar, 200 *µ*m. 3V, third ventricle; dmArc, dorso-medial arcuate nucleus. **(C)** Number of DAT-tdT^+^ and TH^+^ neurons per section in the dmArc (n = 6–7 mice; two-way repeated-measures ANOVA: Age, *F* (6, 36) = 38.84, ^*\*\*\*\**^*p <* 0.0001; Marker, *F* (1, 36) = 126.60, ^*\*\*\*\**^*p <* 0.0001; Age *×* Marker, *F* (6, 36) = 2.70, ^*^*p* = 0.029). **(D)** Percentage of TH^+^ neurons co-labeled for DAT-tdT across postnatal development. **(E)** Representative confocal images of dopaminergic innervation in the ME at P3 (left) and P15 (right). Top, merged DAT-tdT (magenta) and TH (cyan) channels; middle, DAT-tdT channel; bottom, TH channel. The yellow region of interest (ROI) indicates the manually annotated ME. Scale bar, 200 *µ*m. **(F)** Area occupied by DAT-tdT^+^ and TH^+^ neurites in the ME (n = 6–13 sections from 4–7 mice; two-way repeated-measures ANOVA: Age, *F* (4, 43) = 120.50, ^*\*\*\*\**^*p <* 0.0001; Marker, *F* (1, 43) = 115.90, ^*\*\*\*\**^*p <* 0.0001; Age *×* Marker, *F* (4, 43) = 33.96, ^*\*\*\*\**^*p <* 0.0001).

Given that TIDA neurons are the primary regulators of Prl release from the anterior pituitary, we next asked whether anatomical and functional changes in these neurons accompany the rise in circulating Prl. First, we assessed whether the TIDA population changes in number postnatally. To monitor the dopaminergic neurons within the dmArc, we performed immunofluorescence staining of dopamine markers at several postnatal time points. In brain sections obtained from dopamine transporter (DAT)-tdTomato (DAT-tdT) mice,

TIDA neurons were identified by their anatomical location within the Arc and by expression of either tdT or tyrosine hydroxylase (TH), the rate-limiting enzyme for dopamine synthesis. These mice have previously been validated for identification of TIDA neurons (31).

We observed a notable postnatal increase in the number of dopaminergic neurons (**Fig. 1B-C**; main effect of age, ^*\*\*\*\**^*p <* 0.0001), without any significant sex differences (**Fig. S1A-B**; main effect of sex, *p >* 0.05 for DAT-tdT and TH immunoreactive signals). Co-expression analysis revealed that DAT-tdT expression lagged behind TH expression at the earlier time points (postnatal day 1–5; P1–P5), reaching a plateau of ∼80% co-expression by P7 (**Fig. 1D**). This developmental trajectory was paralleled by a reduction in dividing cells in the dmArc, as detected by immunoreactivity for the proliferative marker Ki67 (40, 41) (**Fig. S1C-D**; ^*\*\*\**^*p <* 0.001). Thus, the TIDA neuronal population, as defined by dopaminergic markers, increases considerably to its adult size during the first postnatal weeks, concurrent with a rise in serum Prl (**Fig. 1A**).

To assess if the change in TIDA somata number is paralleled by alterations in the innervation provided by these neurons, we quantified the area covered by DAT-tdT and TH signal at the terminal site in the ME across postnatal development. We were unable to detect significant TH or DAT-tdT fiber innervation in the ME at the earliest postnatal time points analyzed (P1–P5). A marked increase in the area covered by dopaminergic fibers was observed starting at P7 (**Fig. 1E-F**; main effect of age, ^*\*\*\*\**^*p <* 0.0001), suggesting that during the first postnatal days, TIDA neurons may not yet be structurally capable of releasing dopamine into the hypophyseal portal system.

### TIDA active, but not passive, properties undergo postnatal maturation

Having established that the TIDA neuron population expands during early postnatal development, we next determined their electrophysiological properties through whole-cell patch-clamp recordings. Both the median and maximum spontaneous firing rate of TIDA neurons increased significantly across postnatal development (**Fig. 2A-C**; ^*\*\*\*\**^*p <* 0.0001 for both measures). These changes suggest that the rise in firing rate may be supported by developmental adaptations in action potential (AP) generation.

**Figure 2.**
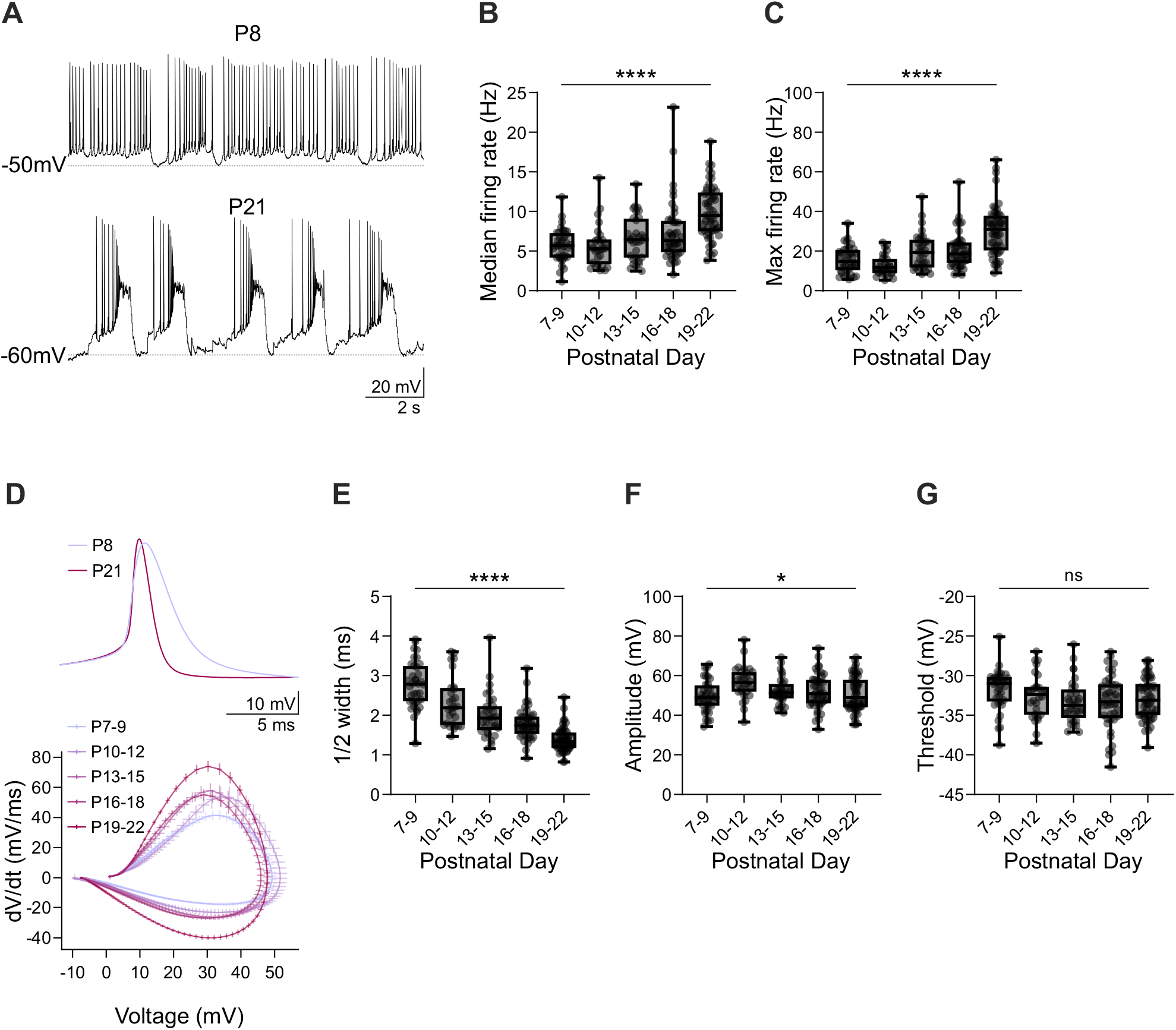
Postnatal maturation of active membrane properties in TIDA neurons. (**A)** Representative whole-cell patch-clamp traces of spontaneous firing activity recorded from TIDA neurons at P8 and P21, note oscillatory firing patterns. **(B)** Box plot showing median firing rate of TIDA neurons across postnatal development (n = 28–54 neurons from 8–15 mice; Kruskal–Wallis test: *H*(4) = 48.95, ^*\*\*\*\**^*p <* 0.0001). **(C)** Box plot showing maximum firing rate of TIDA neurons across postnatal development (n = 28–54 neurons from 8–15 mice; Kruskal–Wallis test: *H*(4) = 61.36, ^*\*\*\*\**^*p <* 0.0001). **(D)** Top, average action potential waveforms recorded from representative neurons at P8 (*n* = 483 APs) and P21 (*n* = 301 APs). Bottom, grand average action potential phase plane plots across postnatal development (n = 28–54 neurons from 8–15 mice). **(E)** Box plot showing action potential half-width across postnatal development (n = 28–54 neurons from 8–15 mice; Kruskal–Wallis test: *H*(4) = 106.6, ^*\*\*\*\**^*p <* 0.0001). **(F)** Box plot showing action potential amplitude across postnatal development (n = 28–54 neurons from 8–15 mice; Kruskal–Wallis test: *H*(4) = 10.20, ^*^*p* = 0.04). **(G)** Box plot showing action potential threshold across postnatal development (n = 28–54 neurons from 8–15 mice; Kruskal–Wallis test: *H*(4) = 4.13, *p* = 0.39).

Accordingly, we examined active membrane properties (**Fig. 2D-G**) and observed a marked decrease in the AP duration at half-amplitude (“half-width”; **Fig. 2D-E**; ^*\*\*\*\**^*p <* 0.0001) and a small, but significant, non-monotonic change in AP amplitude (**Fig. 2F**; ^*^*p <* 0.05). AP threshold, however, remained stable across postnatal development (**Fig. 2G**; *p >* 0.05). Qualitative inspection of the average phase plane plots (**Fig. 2D**) across development confirmed these findings, revealing a progressive narrowing and acceleration of the AP waveform.

In contrast, passive properties such as input resistance and capacitance remained unchanged during the same period (**Fig. S2**; *p >* 0.05 for all measurements). These results indicate that the developmental increase in firing rate may be driven primarily by a shortening of AP duration.

### TIDA oscillations develop postnatally

One hallmark of TIDA neurons is their rhythmic membrane potential oscillations, which vary markedly across reproductive states, sex, and species (31, 42), and are causally linked to circulating Prl levels (18). In male mice, a subpopulation of TIDA neurons exhibits regular oscillations characterized by prominent DOWN states alternating with UP state periods of active firing (31).

To examine how these oscillations develop postnatally, we quantified rhythmic activity by analyzing the autocorrelograms of the membrane potential. Oscillatory strength was estimated by measuring the autocorrelogram peak at positive time lags. To objectively identify neurons with significant rhythmicity, we applied a block bootstrapping-based approach (see **Materials and Methods**). Neurons were classified as “Regular” oscillators when the positive autocorrelogram peak exceeded the 95th percentile of the bootstrap-derived null distribution, and as “Non-regular” firing neurons otherwise (**Fig. 3A-B**).

**Figure 3.**
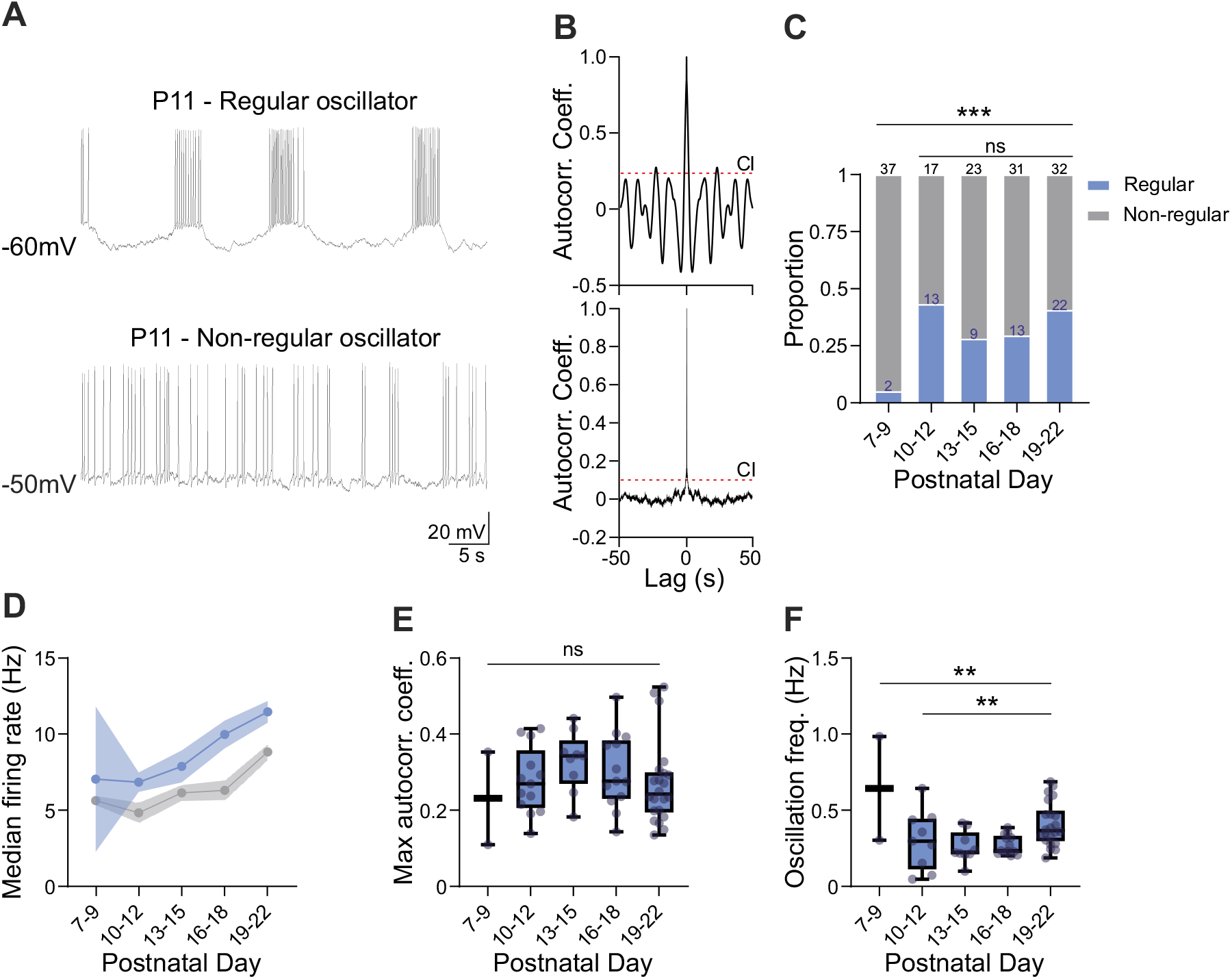
Postnatal maturation of oscillatory properties in TIDA neurons. **(A)** Representative whole-cell patch-clamp traces from two TIDA neurons recorded on postnatal day 11 displaying different types of membrane oscillations. **(B)** Autocorrelogram of a Regular (top) and Non-regular (bottom) oscillating TIDA neuron, corresponding to the examples shown in **(A).** Dashed line indicates the 95% confidence interval (CI). **(C)** Proportion of Regular *vs*. Non-regular neurons across postnatal development (Fisher’s exact test: from P7–9 to P19–22, ^*\*\*\**^*p* = 0.0005; from P10–12 to P19–22, *p* = 0.42; numbers of recorded cells indicated in histogram). **(D)** Line plot showing the average median firing rate (mean *±* s.e.m.) of Regular and Non-regular TIDA neurons across postnatal development (n = 2–34 neurons from 8–15 mice; two-way ANOVA: Age, *F* (4, 184) = 12.07, ^*\*\*\*\**^*p <* 0.0001; Group, *F* (1, 184) = 15.22, ^*\*\*\**^*p* = 0.0001; Age *×* Group, *F* (4, 184) = 0.60, *p* = 0.67). **(E)** Box plot showing the maximum autocorrelation coefficient of Regular TIDA neurons across postnatal development (n = 2–22 neurons from 8–15 mice; Kruskal–Wallis test: *H*(4) = 4.40, *p* = 0.35). **(F)** Box plot showing the oscillation frequency of Regular TIDA neurons across postnatal development (n = 2–22 neurons from 8–15 mice; Kruskal–Wallis test: from P7–9 to P19–22, *H*(4) = 14.99, ^*\*\**^*p* = 0.0047; from P10–12 to P19–22, *H*(3) = 13.19, ^*\*\**^*p* = 0.0043).

Interestingly, we observed an abrupt increase in the number of Regular TIDA neurons between the first and second postnatal time points analyzed, followed by a plateau (**Fig. 3C**; from P7–9 to P19–22: ^*\*\*\**^*p <* 0.001; from P10–12 to P19–22: *p >* 0.05).

To assess whether the previously observed electrophysiological adaptations reflect changes across the entire TIDA population or are specifically driven by the emergence of oscillatory dynamics, we compared neurons grouped by functional class (Regular *vs*. Non-regular). We plotted the median firing rate (MFR) for each group across development (**Fig. 3D**). Regular neurons exhibited significantly higher MFRs compared to Non-regular neurons at all time points (Group effect: ^*\*\*\**^*p <* 0.001), but both groups showed a similar monotonic age-dependent increase (Age effect: ^*\*\*\*\**^*p <* 0.0001; Age *×* Group interaction: *p >* 0.05). This observation suggests that the developmental increase in firing rate occurs independently of the oscillatory behavior, indicating shared underlying mechanisms across functional subtypes.

To further examine how oscillatory properties evolve during postnatal development, we quantified the oscillation strength (**Fig. 3E**) and frequency (**Fig. 3F**) of Regular neurons. While oscillation strength remained stable across age (*p >* 0.05), we observed a small yet significant increase in oscillation frequency (^*\*\**^*p <* 0.01) during later developmental stages.

Our findings suggest that oscillations in TIDA neurons appear abruptly, consistent with a threshold-like transition to rhythmic activity.

### TIDA neuron synaptic properties mature postnatally without affecting the E/I ratio

Having demonstrated that TIDA neurons undergo marked changes during the early postnatal period, we next explored synaptic inputs to these cells. To this end, we recorded spontaneous excitatory and inhibitory postsynaptic currents (sEPSCs and sIPSCs, respectively; **Fig. 4A**).

**Figure 4.**
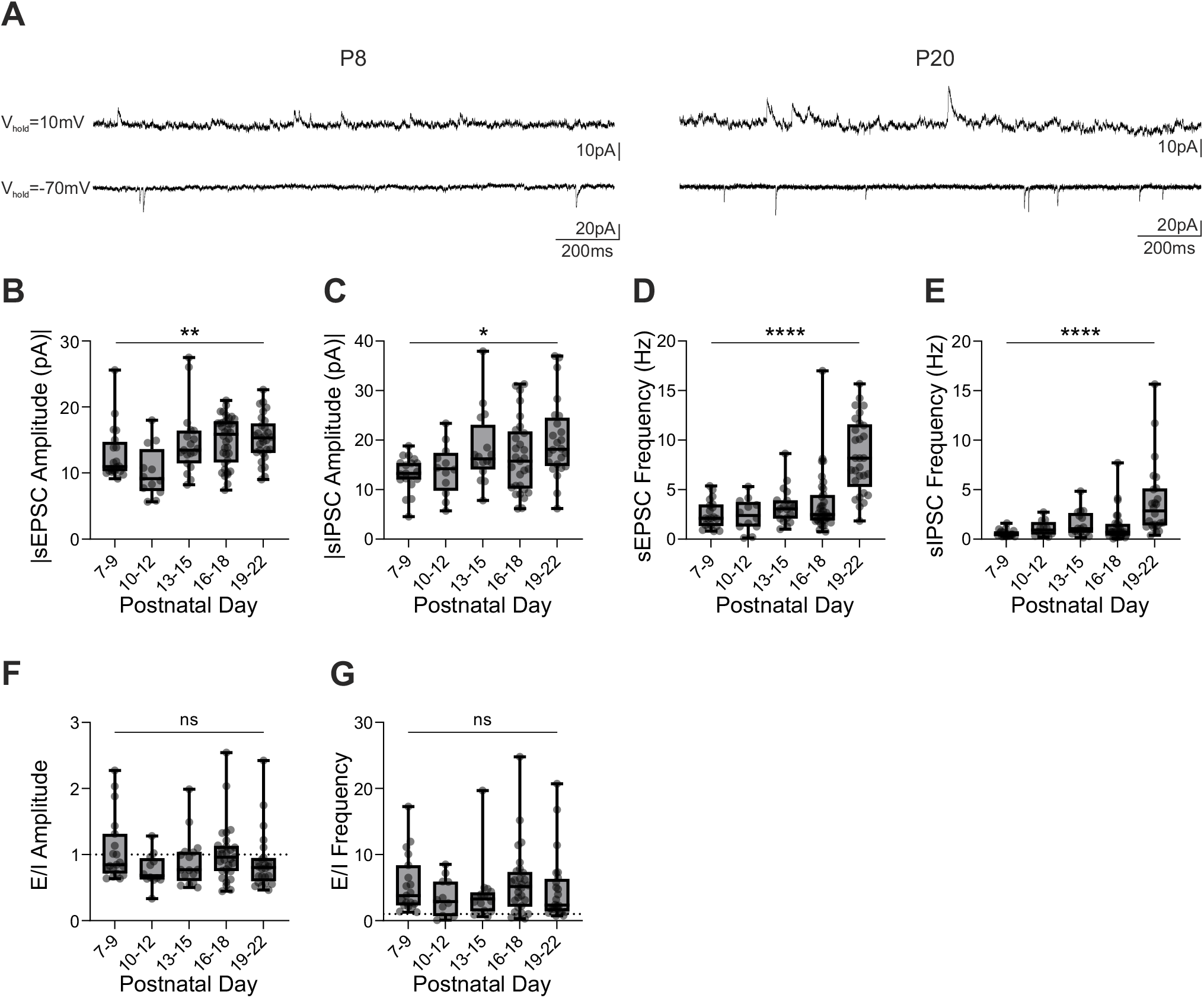
Postnatal maturation of TIDA neurons’ synaptic properties. **(A)** Representative sIPSC (top) and sEPSC (bottom) traces recorded at different holding potentials (*V*_hold_) from TIDA neurons at P8 (left) and P20 (right). Note increases in frequency and amplitude in older animals. **(B)** Box plot showing the average sEPSC amplitude across postnatal development (n = 12–36 neurons from 4–13 mice; Kruskal–Wallis test: *H*(4) = 18.16, ^*\*\**^*p* = 0.0012). **(C)** Box plot showing the average sIPSC amplitude across postnatal development (n = 12–29 neurons from 4–13 mice; Kruskal–Wallis test: *H*(4) = 13.14, ^*^*p* = 0.011). **(D)** Box plot showing the average sEPSC frequency across postnatal development (n = 12–36 neurons from 4–13 mice; Kruskal–Wallis test: *H*(4) = 44.79, ^*\*\*\*\**^*p <* 0.0001). **(E)** Box plot showing the average sIPSC frequency across postnatal development (n = 12–29 neurons from 4–13 mice; Kruskal–Wallis test: *H*(4) = 31.86, ^*\*\*\*\**^*p <* 0.0001). **(F)** Box plot showing the E/I amplitude ratio across postnatal development (n = 12–29 neurons from 4–13 mice; Kruskal–Wallis test: *H*(4) = 6.19, *p* = 0.19). **(G)** Box plot showing the E/I frequency ratio across postnatal development (n = 12–29 neurons from 4–13 mice; Kruskal–Wallis test: *H*(4) = 4.86, *p* = 0.30).

Both sEPSC and sIPSC amplitudes and frequencies increased in an age-dependent manner, although with distinct temporal profiles. The amplitude of both event types showed a gradual and modest increase across development, suggesting progressive but relatively minor maturation of postsynaptic receptor function or subunit composition (**Fig. 4B-C**; sEPSC: ^*\*\**^*p <* 0.01; sIPSC: ^*^*p <* 0.05).

In contrast, the frequency of both sEPSCs and sIPSCs increased dramatically and abruptly between the last two time points analyzed (P16–18 to P19–22), indicating a substantial step-wise increase in presynaptic activity or connectivity during this window (**Fig. 4D-E**; ^*\*\*\*\**^*p <* 0.0001 for both measurements).

To determine whether the observed synaptic maturation affects the balance of incoming excitatory and inhibitory signals during postnatal development, we calculated the excitation/inhibition (E/I) ratio using event amplitude and frequency as proxies for post- and pre-synaptic function, respectively. Notably, neither the E/I amplitude nor the E/I frequency changed significantly over time, indicating that the maturation of synaptic inputs occurs in a coördinated manner that preserves the homeostatic balance between excitation and inhibition at both post- and pre-synaptic levels (**Fig. 4F-G**; *p >* 0.05 for both measurements).

### Principal Component Analysis reveals a monotonic electrophysiological developmental trajectory for TIDA neurons

Next, we sought to identify a developmental trajectory that integrates the functional properties of TIDA neurons across postnatal development.

We selected 98 out of 194 recorded TIDA neurons for which a complete set of electrophysiological features was available and performed principal component analysis (PCA) for dimensionality reduction (passive membrane properties were excluded because none of the extracted parameters showed significant developmental changes). The first two principal components together accounted for 44.24% of the total variance in the dataset.

Neurons appeared to align to an age-related gradient along the first principal component (PC1; **Fig. 5A, S3A**), suggesting a monotonic developmental progression in their electrophysiological profile. As expected, PC1 scores correlated significantly with postnatal age (**Fig. 5B, S3B**; ^*\*\*\*\**^*p <* 0.0001), and we observed significant differences across age groups (**Fig. 5C**; ^*\*\*\*\**^*p <* 0.0001). The electrophysiological parameters most strongly contributing to PC1 included sEPSC and sIPSC frequency and amplitude (**Fig. S3A**), indicating that changes in synaptic activity largely drive the developmental variance captured by this component. PC1, and to a much lesser extent PC8 and 9 (accounting together for only 7.99% of total variance), showed significant correlations with postnatal age (**Fig. S3B**), confirming that PC1 represents the main developmental axis in the dataset.

**Figure 5.**
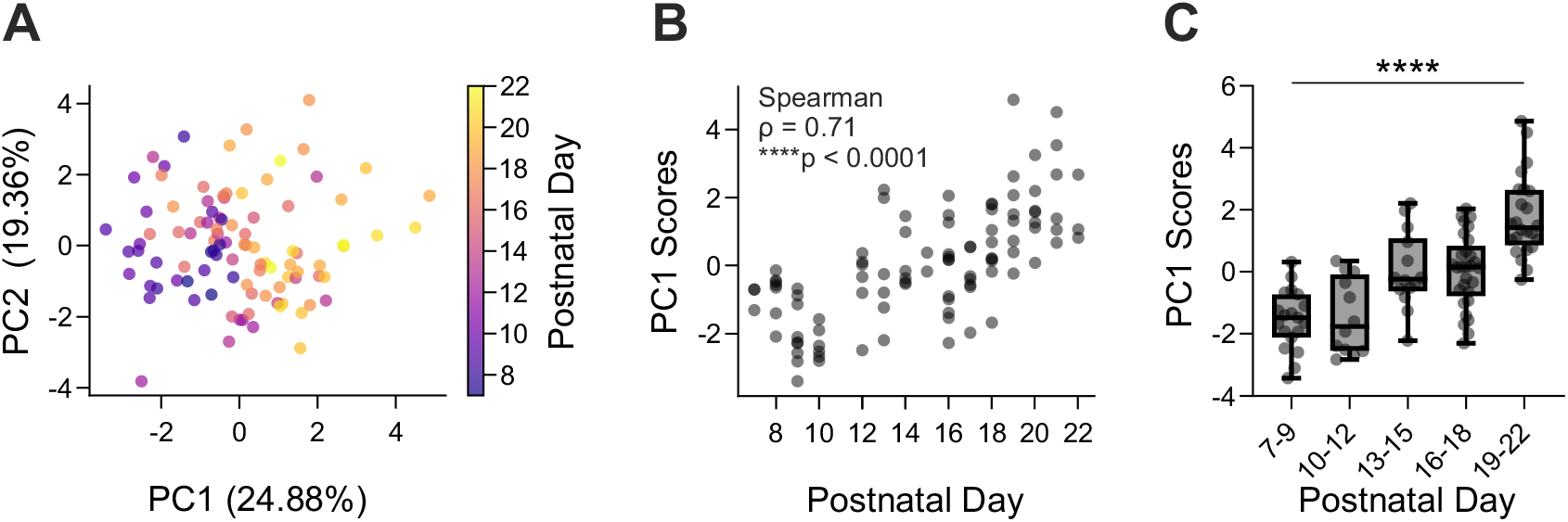
TIDA neurons’ coördinated functional developmental trajectory. **(A)** Principal Component Analysis (PCA) plot showing the distribution of TIDA neuron electrophysiological features, including active membrane properties, firing and oscillatory behavior, and synaptic parameters, across postnatal development. Each dot represents a single recorded neuron, color-coded by age. Principal components 1 and 2 (PC1, PC2) account for 24.88% and 19.36% of the total variance, respectively. **(B)** Spearman correlation between PC1 scores and age (n = 98 neurons from 43 mice; Spearman’s correlation: *ρ* = 0.71, ^*\*\*\*\**^*p <* 0.0001). **(C)** Box plot showing the PC1 scores across postnatal development (n = 12–29 neurons from 4–13 mice; one-way ANOVA: *F* (4, 93) = 23.48, ^*\*\*\*\**^*p <* 0.0001).

This developmental gradient highlights the coördinated maturation of intrinsic and synaptic properties in TIDA neurons, indicating that their functional development is not unconnected or limited to isolated features, but rather follows a structured and coherent program. This trajectory may underlie the gradual establishment of TIDA neurons’ physiological role as the primary regulators of Prl secretion.

### TIDA network functional connections increase during postnatal development, maintaining constant strength

To investigate how TIDA microcircuit connectivity develops over time, we performed wide-field Ca^2+^ imaging in acute brain slices from DAT-GCaMP6s mice. We quantified functional interactions between TIDA neurons located within the same arcuate hemisphere by computing pairwise cross-correlograms of Ca^2+^ activity. To isolate significant interactions, we generated a null distribution of peak cross-correlation values using block-bootstrap resampling of the Ca^2+^ traces, following the same procedure used to assess TIDA oscillatory properties (**Fig. 3**; see **Materials and Methods**). This analysis enabled us to identify neuron pairs exhibiting temporally correlated activity beyond chance levels (**Fig. 6A-C**).

**Figure 6.**
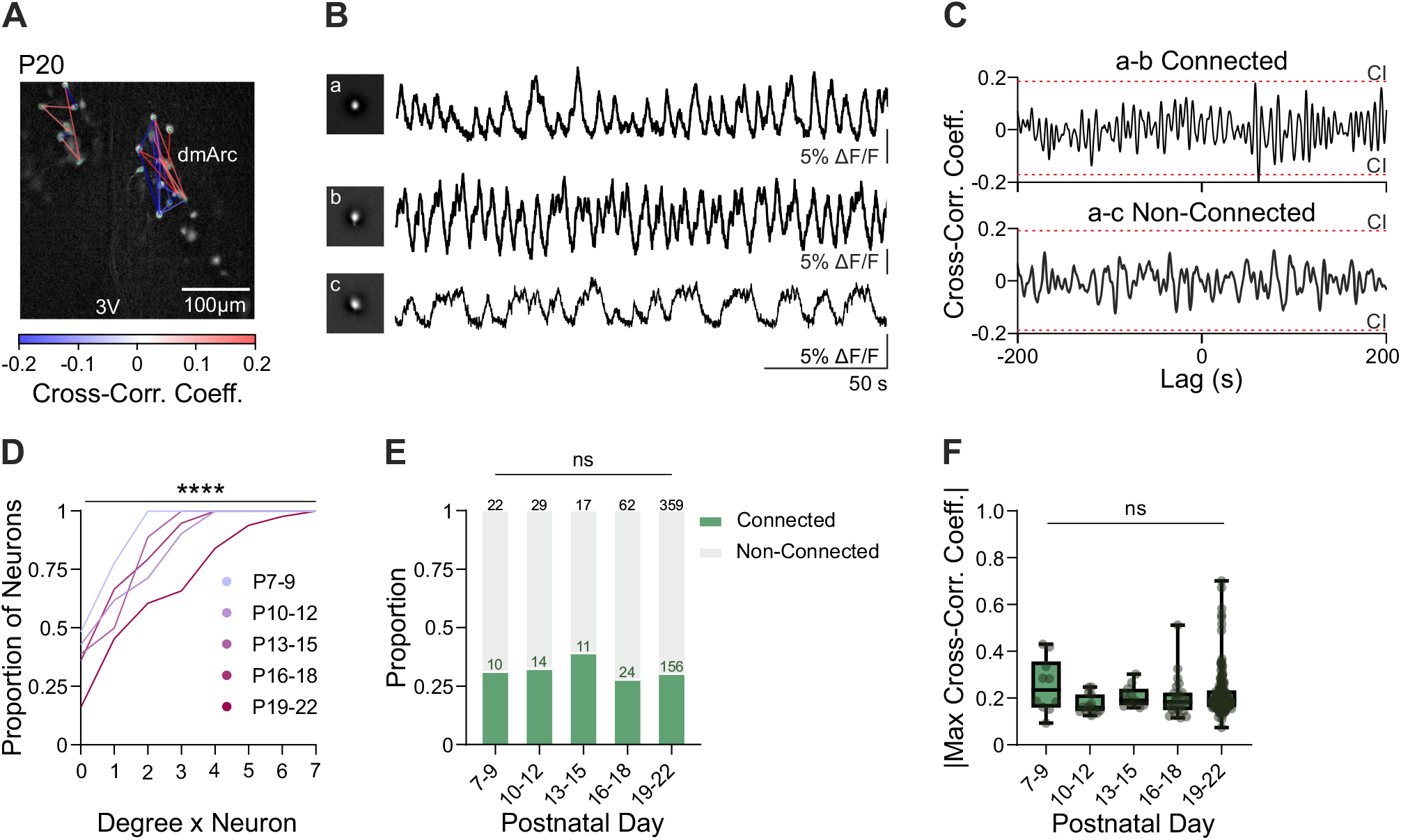
Postnatal refinement of functional connectivity in TIDA neuron networks. **(A)** Representative functional connectivity map of TIDA neurons at P20, superimposed on an *ex vivo* slice preparation. TIDA neurons are marked with green circles; significant pairwise functional connections are represented by lines, color-coded according to their connection strength (cross-correlation coefficient). Scale bar, 100 *µ*m. 3V, third ventricle; dmArc, dorsomedial arcuate nucleus. **(B)** Representative Ca^2+^ traces from three TIDA neurons shown in **(A). (C)** Cross-correlogram computed from the Ca^2+^ traces in **(B)**, showing the temporal correlation between two pairs of neurons. Dashed lines indicate the 5–95% confidence interval used to determine significant connections. **(D)** Cumulative distribution of the number of significant functional connections per neuron (node degree) across postnatal development (n = 18–132 neurons from 2–7 mice; Kruskal–Wallis test: *H*(4) = 28.56, ^*\*\*\*\**^*p <* 0.0001). **(E)** Proportion of connected and non-connected pairs across postnatal development (Fisher’s exact test, *p* = 0.83). **(F)** Box plot showing the absolute value of the maximum cross-correlation coefficient across postnatal development (n = 10–156 pairs from 2–7 mice; Kruskal–Wallis test: *H*(4) = 6.95, *p* = 0.14).

We analyzed the degree (*i.e*., number of functional connections per neuron) from pairwise Ca^2+^ correlation maps. The cumulative degree distribution showed a significant rightward shift with age (**Fig. 6D**; ^*\*\*\*\**^*p <* 0.0001), indicating that individual TIDA neurons become increasingly embedded in the local network. Despite this increase in per-neuron connectivity, the overall proportion of connected neuron pairs remained stable across postnatal ages (**Fig. 6E**; *p >* 0.05), suggesting that the network grows in a coördinated manner without altering its global connection density. Furthermore, the strength of individual connections, as measured by the maximum cross-correlation coefficient, showed no significant developmental trend (**Fig. 6F**; *p >* 0.05), implying that network refinement is primarily driven by changes in connectivity structure rather than connection strength.

### Acute Prl responsiveness is established early in TIDA neurons, whereas basal pathway activation rises postnatally

Finally, in light of the postnatal maturation of TIDA intrinsic, synaptic, and network properties, we examined whether Prl signaling onto TIDA neurons is also dynamic during this period. Mice were acutely injected intraperitoneally with Prl (5 mg/kg) or vehicle (Veh), and activation of the canonical Prl pathway was quantified by immunostaining for phosphorylated Signal Transducer and Activator of Transcription 5 (pSTAT5; 43, 44) in TIDA neurons (**Fig. 7A**). At both postnatal ages examined, Prl administration induced a robust increase in the proportion of pSTAT5positive TIDA neurons compared to Veh controls (P7–9 Veh *vs*. P7–9 Prl: ^*\*\*\*\**^*p <* 0.0001; P19–22 Veh *vs*. P19– 22 Prl: ^*\*\**^*p <* 0.01; **Fig. 7A-B**), indicating that TIDA neurons are Prl-responsive throughout postnatal development. Importantly, the magnitude of Prl-induced pSTAT5 activation did not differ between ages (P7–9 Prl *vs*. P19–22 Prl: *p >* 0.05; **Fig. 7A-B**), suggesting that acute Prl responsiveness is already established early after birth.

**Figure 7.**
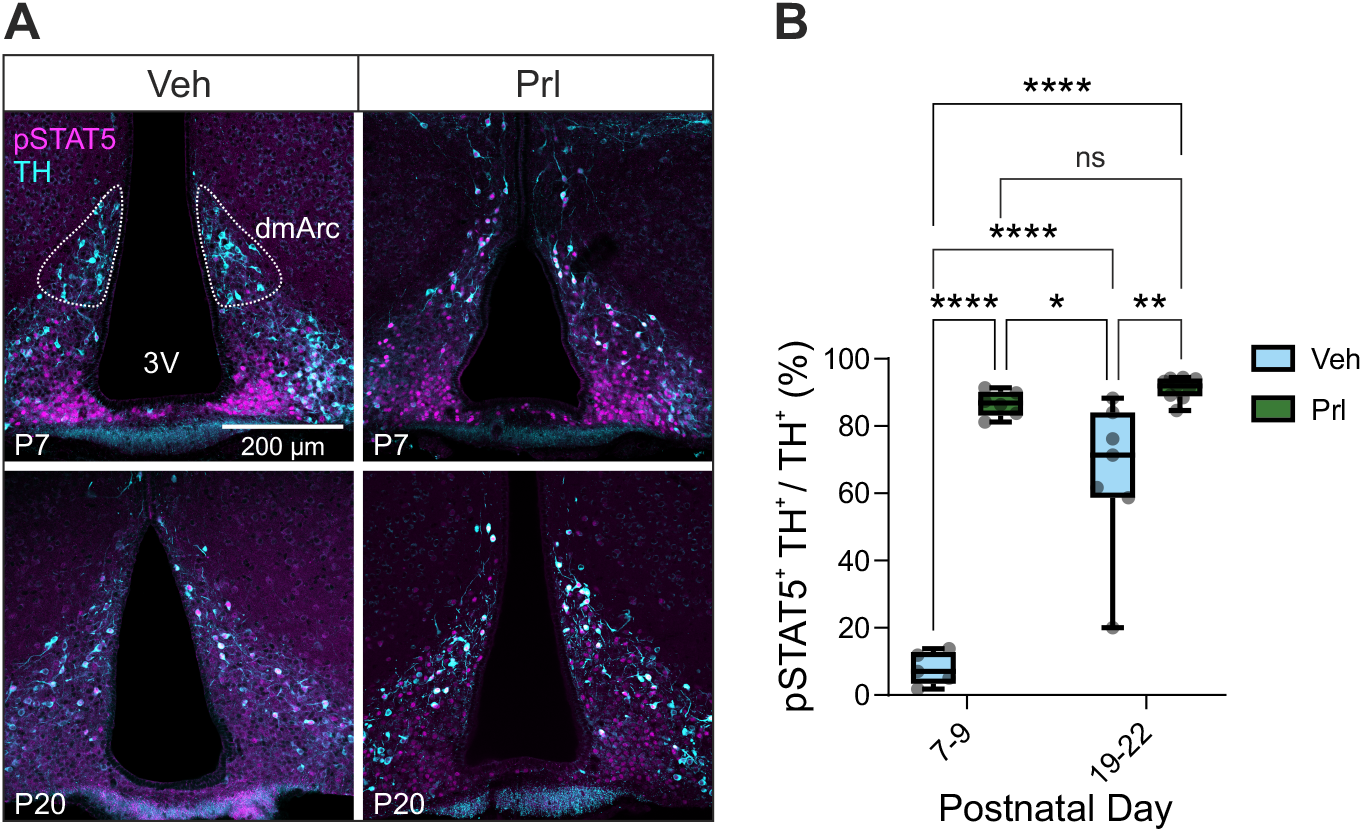
Basal and Prl-induced STAT5 phosphorylation in TIDA neurons during postnatal development. **(A)** Representative confocal images of the mediobasal hypothalamus at P7 (top) and P20 (bottom) following vehicle (Veh; left) or prolactin (Prl; right) administration, showing TIDA neurons identified by immunofluorescence for TH (cyan) and pSTAT5 (magenta) as a readout of Prl signaling. Scale bar, 200 *µ*m. 3V, third ventricle; dmArc, dorsomedial arcuate nucleus. **(B)** Quantification of the proportion of pSTAT5-positive TIDA neurons at P7–9 and P19–22 under Veh and Prl conditions (P7–9 Veh, n = 5 mice; P7–9 Prl, n = 6 mice; P19–22 Veh, n = 7 mice; P19–22 Prl, n = 9 mice). Two-way ANOVA revealed significant main effects of treatment (*F* (1, 23) = 118.50, ^*\*\*\*\**^*p <* 0.0001) and age (*F* (1, 23) = 42.53, ^*\*\*\*\**^*p <* 0.0001), and a significant age *×* treatment interaction (*F* (1, 23) = 30.87, ^*\*\*\*\**^*p <* 0.0001) (post hoc Tukey: P7–9 Veh *vs*. P7–9 Prl, ^*\*\*\*\**^*p <* 0.0001; P19–22 Veh *vs*. P19–22 Prl, ^*\*\**^*p* = 0.002; P7–9 Veh *vs*. P19–22 Veh, ^*\*\*\*\**^*p <* 0.0001; P7– 9 Veh *vs*. P19–22 Prl, ^*\*\*\*\**^*p <* 0.0001; P7–9 Prl *vs*. P19–22 Veh, ^*^*p* = 0.025; P7–9 Prl *vs*. P19–22 Prl, *p* = 0.89).

In contrast, basal pSTAT5 levels measured under Veh conditions showed a significant age-dependent increase (P7–9 Veh *vs*. P19–22 Veh: ^*\*\*\*\**^*p <* 0.0001; **Fig. 7A-B**), indicating higher baseline activation in older animals. Thus, while the capacity of TIDA neurons to respond acutely to circulating Prl is preserved across postnatal development, tonic activation of the STAT5 pathway increases with age. Together, these data reveal a developmental shift in baseline Prl signaling in TIDA neurons, with no change in acute responsiveness.

## Discussion

Postnatal development represents a critical window for the maturation and refinement of neural circuits (see 45), including synaptic pruning and maturation of intrinsic membrane properties (see 46). While the postnatal maturation of cortical and hippocampal circuits has been thoroughly characterized for decades (*e.g*., 47, 48), studies of hypothalamic development, particularly regarding neuroendocrine and homeostatic circuit organization, remain rare, with few exceptions *(e.g*., 35, 36, 49, 50). Notably, the Arc regulates essential physiological processes that emerge or reorganize during early life, such as feeding, metabolic homeostasis, puberty onset, and reproductive control. The maturation of these diverse circuits within a shared anatomical substrate could underscore the importance of the Arc as a key site for coördinated neuroendocrine and homeostatic integration during the postnatal period. Our study focused on the postnatal development of TIDA neurons in the dmArc, whose electrical, cellular, and network properties are unusually well characterized, both in their basal properties (30, 31, 51), feedback control, and their relevance for dopamine release and parental functions (18, 34, 52). However, it has remained unclear how this finely tuned homeostatic circuitry is originally established.

Here, we demonstrate that mouse TIDA neurons undergo a coördinated developmental program during the first three postnatal weeks. This process involves (1) expansion of the TIDA population, (2) a delayed emergence of ME innervation, (3) progressive refinement of intrinsic excitability and rhythmic activity, and (4) a balanced increase in synaptic and local network connectivity. Together, these findings suggest that the transition of the TIDA-Prl axis from an immature to a functionally competent state is governed by a structured developmental sequence that supports the gradual establishment of neuroendocrine homeostasis.

We found that the number of TIDA neurons increases significantly during early postnatal development. While our approach did not allow us to directly determine whether this rise results from postnatal neurogenesis, originating from previously identified hypothalamic progenitor niches in the Arc (53, 54), or from the final stages of dopaminergic differentiation (*e.g*., acquisition of TH expression), our data show that the increase slows down markedly after P7 (**Fig. 1C**). This temporal pattern suggests that the structural establishment of the TIDA precedes the full maturation of the Prl secretory system, which continues to expand, through lactotroph differentiation and progressive acquisition of adult-like Prl secretion, over the first eight postnatal weeks (20, 21).

The relatively late acquisition of dopaminergic markers in the dmArc, in agreement with findings in the rat (55), contrasts with the earlier specification of midbrain dopaminergic neurons, which already express TH by E11.5 (56). This temporal distinction likely highlights region-specific developmental trajectories of dopaminergic differentiation, reflecting divergent regulatory programs that govern hypothalamic versus mesencephalic dopaminergic neuronal maturation. Our findings indicate that the functional capacity of TIDA neurons to regulate Prl secretion is progressively acquired during early postnatal development. This is supported by both the expansion of the TIDA population described above and multiple converging lines of evidence.

First, dopaminergic markers within the ME are virtually undetectable before postnatal days 5–7 (**Fig. 1E-F**), suggesting a delayed arrival or maturation of TIDA axons within the ME. This delayed innervation makes TIDA neurons unlikely to release significant amounts of dopamine in the hypophyseal portal system during the first week after birth. Consistent with this, the ability of haloperidol, a dopamine D2 receptor antagonist, to elevate serum Prl in rats is only acquired several days after birth, increasing markedly with age (57, 58). Second, TIDA neurons exhibit a gradual maturation of their intrinsic electrophysiological properties. During the postnatal period, they increase their ability to generate spontaneous, sustained, high-frequency spike trains (**Fig. 2**), consistent with developmental trajectories observed in other neuronal populations (59–61). This increase in firing capacity likely enhances their ability to reliably release dopamine, contributing to a strengthened restraint of Prl secretion. Notably, late in postnatal development (P19–22), firing frequency has reached the level previously identified to maximize dopamine yield (**Fig. 2B**; 34). Third, regular membrane potential oscillations, a feature functionally linked with TIDA neuron inhibition of Prl (18), emerged only from P10–12 (at earlier timepoints, such patterns were rare: 2/39 recorded neurons; **Fig. 3**). Given the importance of TIDA oscillations in Prl regulation, the sharp postnatal increase in regular oscillators likely marks an important developmental step in the maturation of negative feedback control over Prl.

Fourth, TIDA neurons exhibited a marked postnatal increase in sEPSC and sIPSC amplitude and frequency (**Fig. 4**). This parallel enhancement of excitatory and inhibitory transmission suggests an expanding capacity of TIDA neurons to integrate afferent neural signals. Notably, despite the overall increase in synaptic drive, the E/I ratio remained remarkably stable across development. While postnatal maturation in many brain regions is characterized by pronounced shifts in E/I balance (62, 63), its preservation in TIDA neurons may indicate a developmental trajectory optimized for reliable signal integration.

Lastly, TIDA neurons progressively consolidate their integration within the local network (**Fig. 6**). The augmented per-neuron connectivity likely stems, at least in part, from the concurrent increase of the TIDA neuron population and incorporation into the microcircuit. Importantly, TIDA network expansion occurs without detectable changes in overall network density or connection strength, suggesting that circuit growth follows a structured scaling rule rather than an indiscriminate increase of functional interactions. Such an organization may preserve stability and information flow, reflecting the same principle of balanced scaling that governs synaptic maturation at the single-cell level.

Taken together, the progressive emergence of mature firing capacity, rhythmic activity, and network integration indicates that TIDA neurons transition from an immature to a functionally competent state through a coördinated developmental process, rather than through the independent acquisition of isolated electrophysiological features. Consistent with this view, multivariate analysis revealed a clear age-dependent organization of intrinsic and activity-related properties along a continuous trajectory (**Fig. 5, S3**), capturing the gradual consolidation of functional firing patterns across postnatal development. This monotonic progression supports the idea that postnatal maturation of TIDA neurons reflects a structured and unified process aligned with their emerging role in neuroendocrine regulation.

Although our data would suggest an age-dependent enhancement of the inhibitory drive exerted by TIDA on Prl, we observed a significant postnatal increase in serum Prl levels, consistent with previous studies (**Fig. 1A**; 23–25 This seemingly paradoxical rise may reflect the concurrent increase in pituitary Prl production capacity, driven by both an upregulation of Prl mRNA expression and an expansion of the lactotroph population during this period (20– In this view, the postnatal rise in circulating Prl is not a consequence of failed or delayed inhibitory control, but rather reflects a system in which the rate of lactotroph expansion and the concurrent increase in secretory output exceed the capacity of the maturing TIDA inhibitory tone to fully constrain it, resulting in a net rise in circulating Prl despite the progressive consolidation of dopaminergic control. Thus, the net result is a controlled postnatal rise in circulating Prl toward resting adult levels, driven by lactotroph expansion rather than a failure of dopaminergic inhibition. In general terms, it may be beneficial to allow the establishment of a system before its inhibitory feedback control is fully in place. A corollary is found in the postnatal maturation of cortical circuits, where excitatory networks become active before the GABAergic inhibitory interneurons that will ultimately control them reach functional maturity and synaptic integration, a developmental lag considered to gate the onset of critical period plasticity (64).

Within this permissive developmental regime, Prl signaling in early postnatal stages likely contributes to the coördinated maturation of the TIDA-Prl axis. TIDA neurons respond robustly to acute Prl stimulation from early postnatal stages, whereas basal pSTAT5 signaling increases with age (**Fig. 7**). This pattern suggests that the molecular machinery for Prl detection is established early, while tonic exposure rises as lactotroph output increases. Such a developmental trajectory aligns with findings in Prl-deficient mice, where early postnatal Prl administration promoted TIDA neuron differentiation (65), highlighting a critical window during which Prl acts as a neurotrophic factor. Together, these observations are consistent with a model in which the TIDA-Prl axis is shaped by a positive developmental cascade: expanding lactotroph output drives rising Prl levels, which in turn promote TIDA maturation, likely in concert with intrinsic developmental programs, ultimately consolidating the inhibitory tone that stabilizes Prl at adult levels.

This scenario may reflect a general principle of developmental regulation, in which both the effector (lactotroph output) and its controller (TIDA-mediated inhibition) mature progressively and in coördination during the postnatal period (**Fig. 8**). This structured maturation may allow the system to transition into an adult configuration while maintaining stability. Postnatal maturation of the TIDA-Prl axis would then exemplify how neuroendocrine function emerges from the consolidation of circuit elements that interact in concert, rather than from the independent maturation of individual components, thereby optimizing hormonal output and regulatory control in parallel. Circulating hormones such as Prl itself and growth hormone could serve as molecular guides for this process, as both have been shown to influence TIDA neuron differentiation and maintenance (*e.g*., 65, 66). *Future work* should dissect how these signaling pathways drive each maturation step, and explore how perturbations, such as peri-natal stress or pharmacological insults *(e.g*., antipsychotics and SSRIs, known to affect the TIDA-Prl axis *(e.g*., 66–69)), impact the developmental trajectory and ultimately reproductive and parental behaviors later in life.

**Figure 8.**
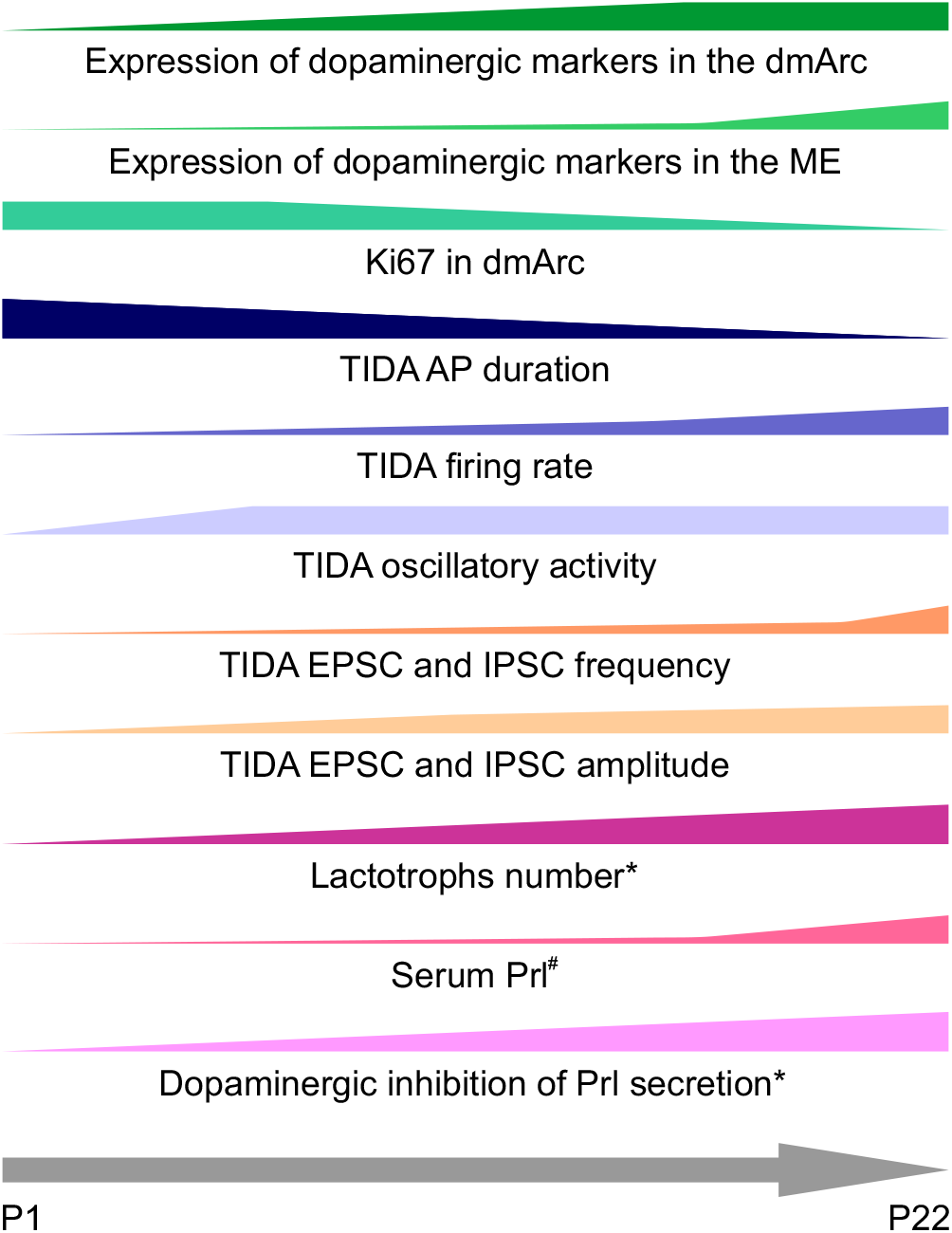
Coördinated postnatal maturation of the TIDA-Prl axis. Schematic summary of the molecular, electrophysiological, and endocrine changes occurring between P1 and P22. Wedges represent the relative increase or decrease of each parameter across postnatal development and are intended for qualitative illustration only. Asterisks (*) denote findings previously reported by other groups, including the postnatal increase in lactotroph number (20, 21) and the age-dependent acquisition of haloperidol-induced increases in circulating Prl in rats (57, 58). Hashtags (#) denote findings previously reported (23–25) and independently confirmed in the present study.

## Materials and Methods

### Animals

All experiments were approved by the local ethical committee (*Stockholms Norra Djurförsöksetiska Nämnd*). Procedures adhered to the European Parliament and Council of the European Union Directive (2010/63/EU) of September 22, 2010. Mice of both sexes, aged P1 to P22, were used for all experiments. DAT-Cre (70), DAT-tdTomato mice generated by crossing DAT-Cre mice with Ai14 reporter mice (B6.Cg-Gt(ROSA)^26Sortm14(CAG-tdTomato)Hze^J; RRID: IMSR_JAX: 007914), and DAT-GCaMP6s mice generated by crossing DAT-Cre mice with Ai96 reporter mice (B6J.Cg-Gt(ROSA)^26Sortm96(CAG-GCaMP6s)Hze^ /MwarJ; RRID: IMSR_JAX: 028866) were maintained on a C57BL/6J background. Animals were housed under a 12 h/12 h light/dark cycle in a temperature- and humidity-controlled environment with *ad libitum* access to food and water. Experimental pups remained with the dam until the day of the terminal experimental procedure.

### Prolactin ELISA

Mice were anesthetized with isoflurane, and trunk blood samples were collected after decapitation. The serum was obtained by leaving the blood to clot at RT for 30 min to 1 h and centrifugation at 4 ^◦^C for 15 min at 2000 *× g*. The supernatant was aliquoted and stored at -80 ^◦^C. Prl serum levels were measured using an ELISA ready-to-use kit (Merck RAB0408). Serum samples were diluted 1:10 in assay buffer. Absorbance at 450 nm was measured in technical duplicates. Average absorbance values were used to calculate concentrations using a 4-parameter logistic (4PL) standard curve.

### Prolactin administration

Mice at P7–9 or P19–22 received intraperitoneal (i.p.) injections of 5 mg/kg recombinant mouse Prl (ProSpec - CYT-321) or an equivalent volume of vehicle (Phosphate Buffer Saline; PBS). Animals were returned to their home cage and perfused (*vide infra*) 45 min post-injection.

### Immunofluorescence

Mice were deeply anesthetized with sodium pentobarbital and intracardially perfused with pre-warmed (37 ^◦^C) Ca^2+^-free Tyrode’s solution, followed by 4 ^◦^C Zamboni’s fixative (4% PFA and 14% saturated picric acid in 0.1 M Millonig Buffer). Whole brains were dissected and immersed in Zamboni’s fixative overnight, and were then transferred to cryoprotectant solution (30% Ethylene glycol, 30% Glycerol in 0.02 M Phosphate Buffer solution) and stored for up to four weeks at -20 ^◦^C.

Coronal brain sections (70 *µ*m thick) were obtained using a vibratome (Campden Instruments, 5100mz) and processed for immunofluorescence (IF). Briefly, free-floating sections were washed twice for 10 min in 0.01 M PBS, then incubated overnight at 4 ^◦^C with primary antibodies diluted in PBS containing 0.3% Triton-X100 and 1% BSA. After washing, sections were incubated for 2 h at RT with fluorophore-conjugated secondary antibodies and counterstained with DAPI (1:10000; Thermo Fisher) for ten minutes. All tissue sections were mounted on glass slides and cover-slipped using Fluoromount-G™ mounting medium (Invitrogen - 00-4958-02).

For pSTAT5 staining, an antigen retrieval step was performed before primary antibody staining by incubating sections in sodium citrate buffer (pH 6.0) at 95 ^◦^C for 15 min, followed by the protocol described above.

The following antibody combinations were used: IF for DAT-tdT and TH: mouse Anti-DsRed-Alexa594 1:500 (Santa Cruz Biotechnology - sc390909AF594) and rabbit Anti-TH 1:1000 (Encor - RPCA-TH) with donkey Anti-Rabbit-Alexa488 1:500 (Invitrogen - A21206). IF for Ki67 and TH: rat Anti-Ki67-Alexa488 1:500 (Invitrogen - 53-5698-80) and rabbit Anti-TH 1:1000 (Encor - RPCA-TH) with donkey Anti-Rabbit-Alexa594 1:500 (Invitrogen - A21207). IF for pSTAT5 and TH: rabbit Anti-pSTAT5 1:100 (Cell Signaling Technologies - 9359) and chicken Anti-TH 1:500 (Chemicon - AB9702) with donkey Anti-Rabbit-Alexa488 (Invitrogen - A21206) and goat Anti-Chicken-Alexa594 1:500 (Invitrogen - A11042), respectively. Fluorescence confocal images were acquired with a Leica STELLARIS 5 microscope.

### Confocal image analysis

Image analysis was performed on Fiji (71). DAT-tdT^+^, TH^+^, and Ki67^+^ cells in the dmArc were counted manually in both hemispheres. To quantify the density of dopaminergic neurites in the ME, images were automatically thresholded using Yen’s method (72), and the percentage of above-threshold pixels was calculated within the manually delineated ME region.

### Slice preparation for electrophysiology

Mice were anesthetized with isoflurane and decapitated, and the brain was rapidly removed and placed in ice-cold sucrose-based slicing solution containing (in mM): sucrose (213.3), KCl (2.0), NaH_2_PO_4_ (1.2), NaHCO_3_ (26.0), glucose (10.0), CaCl_2_ (2.0), MgCl_2_ (2.0), bubbled with 95% O_2_ / 5% CO_2_. Coronal brain slices (250 *µ*m thick) were obtained using a vibratome (Campden Instruments, 7000smz-2), then transferred to an artificial cerebrospinal fluid (aCSF) solution containing (in mM): NaCl (127.0), KCl (2.0), NaH_2_PO_4_ (1.2), NaHCO_3_ (26.0), glucose (10.0), CaCl_2_ (2.4), MgSO_4_ (1.3), bubbled with 95% O_2_ / 5% CO_2_. Slices were incubated at 34 ^◦^C for 15 min and subsequently maintained at RT until use. During recordings, slices were continuously superfused with aCSF at 2 mL/min and maintained at near-physiological temperature (32–35 ^◦^C).

### *Ex vivo* electrophysiological recordings

Whole-cell patch-clamp recordings were performed on DAT-tdT^+^ neurons of the dmArc. Patch pipettes (7– 10 MΩ) were filled with an internal solution containing (in mM): K-gluconate (140.0), KCl (10.0), HEPES (10.0), EGTA (1.0), Na_2_-ATP (2.0), Neurobiotin (6.2), pH adjusted to 7.2–7.3 with KOH, 280–290 mOsm/kg. Data are reported without corrections for the liquid junction potential (+16 mV). Data were acquired with a Multiclamp 700B amplifier and pClamp 11 software (Molecular Devices), filtered at 2 kHz, and digitized at 20 kHz using a Digidata 1440A (Molecular Devices).

### Electrophysiology analysis

Active and passive membrane properties were extracted using Clampfit 11 (Molecular Devices).

Neurons were classified as Regular or Non-regular based on autocorrelogram analysis of 100 s of spontaneous activity. The maximal autocorrelation coefficient at lag > 0 s was compared to a null distribution generated by simple block bootstrapping (1000 iterations; block size 2 s). Neurons exceeding the 95th percentile of the null distribution were classified as Regular.

Spontaneous EPSCs and IPSCs were detected using miniML (73). A single pretrained model was independently fine-tuned for sEPSC and sIPSC detection using manually annotated datasets containing equal proportions of true events and noise, with an 80/20 train-test split (sEPSCs: 890 events from 10 traces; sIPSCs: 960 events from 20 traces). The resulting models achieved classification accuracies of 0.99 (sEPSCs) and 0.97 (sIPSCs).

### Widefield Ca^2+^ imaging recordings

Ca^2+^ imaging was performed under the same perfusion rate and temperature conditions as electrophysiological recordings. Fluorescence images (1024 *×* 1024 pixels) were acquired with a CMOS camera (Prime BSI Express, Teledyne Technologies) at 20 Hz for 600 s controlled by Micro-Manager (74) using a water-immersion objective (W Plan-Apochromat 20x/1.0 NA, Zeiss).

### Ca^2+^ imaging analysis

Ca^2+^ traces were extracted using Inscopix Data Processing Software (Inscopix). Briefly, videos were temporally downsampled by a factor of 2, preprocessed using a spatial bandpass filter (0.001–0.5 pixel^−1^), and motion-corrected. Δ*F/F* was then computed on a per-pixel basis from the motion-corrected video as:

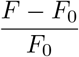

where *F*_0_ corresponds to the mean fluorescence intensity over time for each pixel. TIDA neurons were subsequently identified and manually segmented based on their location within the recorded field of view, and their Ca^2+^ traces were extracted.

Functional connectivity network graphs were built using a custom-written MATLAB script. Pairwise cross-correlograms were computed between Ca^2+^ traces of neurons within the same hemisphere. Connection strength was quantified as the peak with the largest absolute correlation coefficient, representing the maximal correlation magnitude, while retaining the original sign at the peak to preserve the directionality of the interaction. Statistical significance was assessed using simple block bootstrapping (1000 iterations; block size 5 s). Connections falling outside the 5th–95th percentile of the null distribution were considered significant.

### Statistical analysis and data visualizations

Statistical analyses were performed using GraphPad Prism 10 and MATLAB 2022. Sample sizes were chosen to be comparable to those used in similar studies. For all tests, a significance threshold of *α* = 0.05 was applied. Parametric tests were used to evaluate the null hypothesis (no difference among groups) when data met the assumptions of normality, assessed by D’Agostino-Pearson or Shapiro-Wilk tests. When data did not satisfy normality assumptions, appropriate nonparametric tests were applied. Contingency tables were analyzed using Fisher’s exact test. Comparisons involving more than two groups for one independent variable were performed using one-way ANOVA or the Kruskal-Wallis test as the non-parametric alternative. For experiments with more than one independent variable, two-way ANOVA was used. Correlations between variables were assessed using Spearman’s rank correlation. Detailed statistical information is provided in the Results section, figures, and figure legends.

Unless otherwise specified, box plots display the interquartile range with the median indicated by a horizontal line inside the box. Whiskers represent the full data range, and individual data points are overlaid as circles. Statistical significance is indicated as: *ns*, not significant; *p >* 0.05; ^*^*p <* 0.05; ^*\*\**^*p <* 0.01; ^*\*\*\**^*p <* 0.001; ^*\*\*\*\**^*p <* 0.0001.

## Acknowledgments

This work was generously supported by a European Research Council Advanced Grant under the European Union’s Horizon 2020 research and Innovation Programme (TOGETHER; Grant Agreement No. 101021496), the Distinguished Professor program of the Swedish Research Council (Vetenskapsrådet grant 2021-00671) to C.B., and a 2025 Brain and Behavior Research Foundation (BBRF) Young Investigator Grant to A.L. The authors thank Dr. Paul Williams for overseeing the animal colonies, and acknowledge the expert services provided by the Experimental Core Facility (ECF) at Stockholm University, Sweden. The authors express their gratitude to all members of the Broberger laboratory for helpful discussions and valuable experimental input during the execution of this study and preparation of the manuscript.

## Author contributions

A.L.: Conceptualization, Methodology, Investigation, Software, Formal Analysis, Data Curation, Visualization, Writing—Original Draft, Writing—Review & Editing, Funding Acquisition. A.M.: Investigation, Methodology, Formal Analysis, Writing—Review & Editing. J.F.: Conceptualization, Resources, Writing—Review & Editing. C.B.: Conceptualization, Resources, Writing—Review & Editing, Supervision, Project Administration, Funding Acquisition.

## Data, Materials, and Software Availability

Source data for all figures and the original code used in this study are available from the corresponding authors upon request.

