## Supplemental Information for "Stable Network Homeostasis during Multi-Level Postnatal Maturation of the Mouse Tuberoinfundibular Dopamine–Prolactin Axis"

### This PDF file includes:

Figures S1–S3

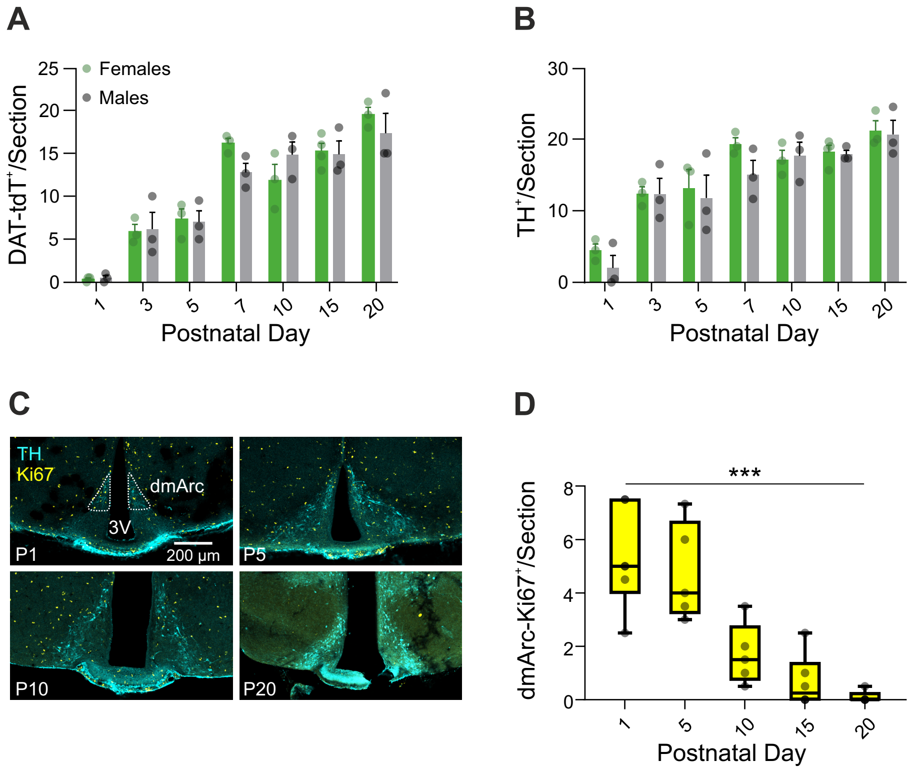


**Figure S1 related to Fig. 1. (A)** Bar graph showing the average number of DAT-tdT$^{+}$ dmArc neurons per section across postnatal development, separated by sex (n = 3–4 mice; two-way ANOVA: Age, $F(6, 29) = 46.06$, $^{****}p < 0.0001$; Sex, $F(1, 29) = 0.33$, $p = 0.57$; Age × Sex, $F(6, 29) = 1.16$, $p = 0.36$). **(B)** Bar graph showing the average number of TH$^{+}$ dmArc neurons per section across postnatal development, separated by sex (n = 3–4 mice; two-way ANOVA: Age, $F(6, 29) = 22.37$, $^{****}p < 0.0001$; Sex, $F(1, 29) = 1.76$, $p = 0.19$; Age × Sex, $F(6, 29) = 0.46$, $p = 0.83$). **(C)** Representative confocal images of the mediobasal hypothalamus at different postnatal days showing dopaminergic neurons identified by TH immunofluorescence (cyan) and proliferating cells labeled with Ki67 (yellow). Scale bar = 200 µm. 3V, third ventricle; dmArc, dorsomedial arcuate nucleus. **(D)** Box plot showing the average number of Ki67$^{+}$ cells per section in the dmArc across postnatal development (n = 5–6 mice; Kruskal–Wallis test: $H(4) = 20.85$, $^{***}p = 0.0003$). **(A,B)** Bars represent the mean; whiskers indicate s.e.m.; individual data points are shown as circles.

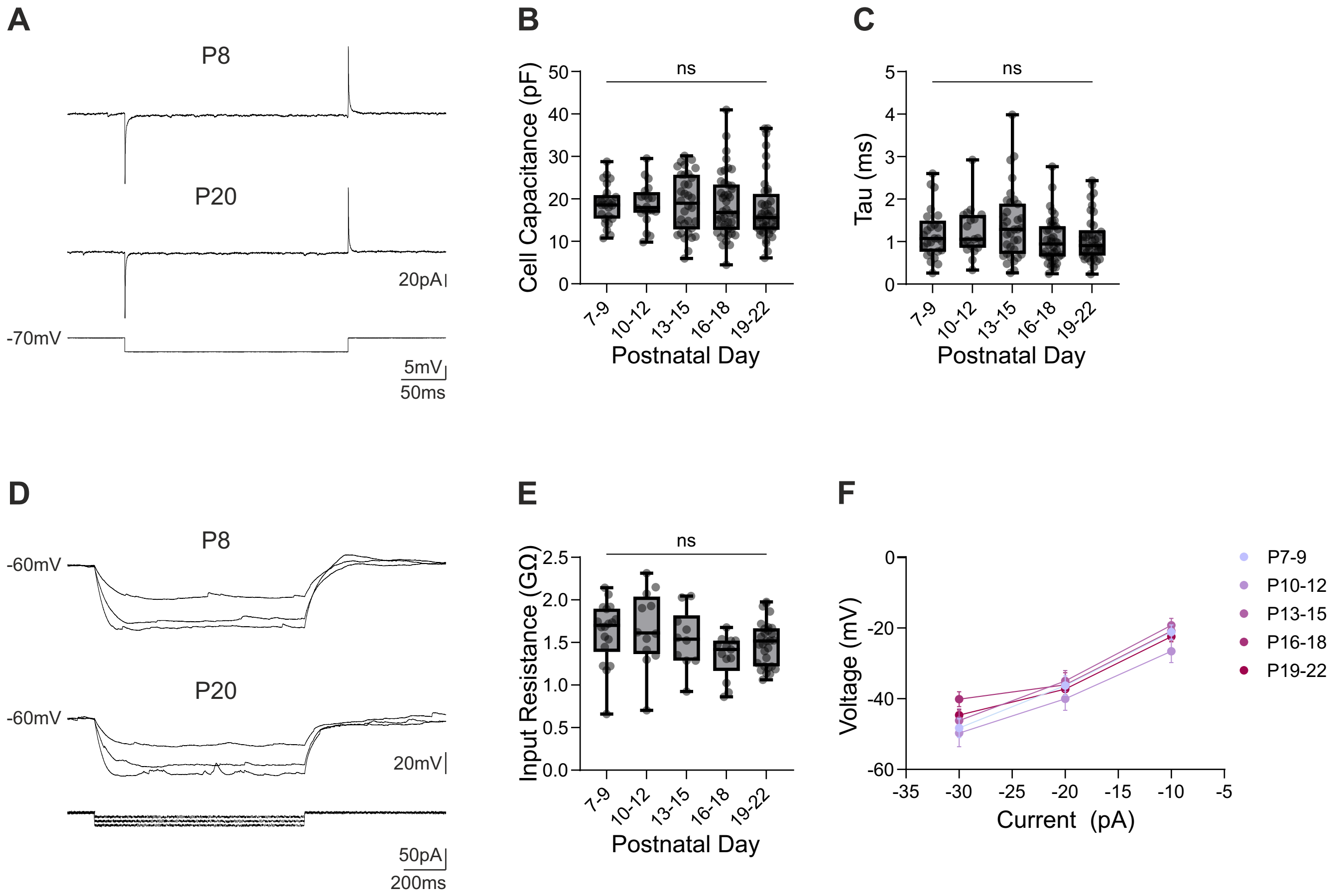


**S2 related to Fig. 2.** **(A)** (Top-middle) Representative whole-cell patch-clamp traces recorded in voltage-clamp mode used to extract membrane capacitance and tau. (Bottom) Corresponding voltage command (from -70 mV to -75 mV). **(B)** Box plot showing TIDA neuron membrane capacitance across postnatal development (n = 20–42 neurons from 8–15 mice; Kruskal–Wallis test: $H(4) = 2.54$, $p = 0.64$). **(C)** Box plot showing TIDA neuron membrane time constant (tau) across postnatal development (n = 20–42 neurons from 8–15 mice; Kruskal–Wallis test: $H(4) = 3.93$, $p = 0.42$). **(D)** (Top-middle) Representative whole-cell patch-clamp traces recorded in current-clamp mode used to extract input resistance. (Bottom) Corresponding injected current steps (from -30 pA to -10 pA in 10 pA steps). **(E)** Box plot showing TIDA neuron input resistance across postnatal development recorded holding neurons at -60 mV (n = 10–29 neurons from 8–15 mice; one-way ANOVA: $F(4, 77) = 2.01$, $p = 0.10$). **(F)** Line plot showing the membrane potential response of TIDA neurons to negative current injections (-30 pA to -10 pA), across postnatal development (n = 10–29 neurons from 8–15 mice). Data are reported as mean ± s.e.m.

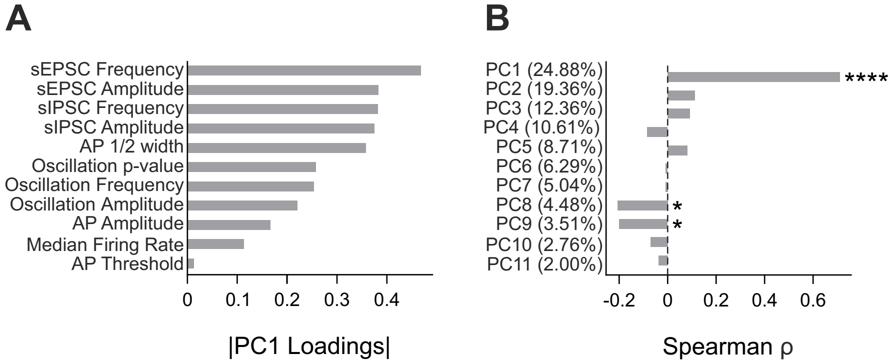


**S3 related to Fig. 5. (A)** Absolute loadings of electrophysiological features on the first principal component (PC1). Active, synaptic, and oscillatory properties were included (passive properties were excluded, as none showed developmental changes). Synaptic features contributed most strongly to PC1. **(B)** Spearman correlation coefficients ($\rho$) between postnatal age and the scores of each principal component. The percentage of total variance explained by each component is indicated in parentheses.
